# A Hypothalamic Inhibitory Circuit Encoding the Scalability of Stress Responses

**DOI:** 10.64898/2026.06.30.735593

**Authors:** Yuhan Cao, Megumi H. Seese, Zhiying Jiang, Cunjin Su, Maojie Yang, Fabricio H. Do Monte, Qingchun Tong, Yuanzhong Xu

**Affiliations:** Brown Foundation Institute of Molecular Medicine, University of Texas Health Science Center at Houston (UTHealth), Houston, TX 77030; MD Anderson Cancer Center & UTHealth Houston Graduate School for Biomedical Sciences, UTHealth; Department of Neurobiology and Anatomy of McGovern Medical School, UTHealth, Houston, TX 77030

**Keywords:** CRH, PVH, Arcuate nucleus, GABAergic, Stress, Scalability

## Abstract

An appropriate stress response is essential for properly responding to, coping with, and subsequently recovering from disturbing environmental stimuli. However, how the brain dynamically encodes the scalability of stress responses remains poorly understood. Here, we found that, GABAergic neurons in the arcuate nucleus (Arc, denoted as Arc^GABA^ neurons) send direct inputs to corticotropin-releasing hormone (CRH) neurons in the paraventricular nucleus of the hypothalamus (PVH, denoted as PVH^CRH^ neurons), the primary regulators of the hypothalamic-pituitary-adrenal (HPA) axis. Although PVH^CRH^ neurons exhibited time-locked activation in response to various environmental stressors, both GABA release onto PVH^CRH^ neurons and the activity of PVH^CRH^-projecting Arc^GABA^ neurons were selectively reduced during exposure to prolonged, high-intensity stressors, but not following exposure to transient, low-intensity stressors. Notably, GABA release onto PVH^CRH^ neurons was positively correlated with PVH^CRH^-projecting Arc^GABA^ neuron activity, yet anticorrelated with PVH^CRH^ neuronal activity in response to the same prolonged, high-intensity stressors. Selective silencing of PVH^CRH-^projecting Arc^GABA^ neurons was sufficient to elevate HPA axis activity and stress levels, phenocopying the effect of direct of PVH^CRH^ neuron activation. Conversely, selective activation of PVH^CRH-^projecting Arc^GABA^ neurons reduced both HPA axis activity and stress levels, this effect was completely abolished by concurrent excitation of PVH^CRH^ neurons. Molecular identity screening further revealed that these PVH^CRH^-projecting Arc^GABA^ neurons are not subsets expressing agouti-related peptide (AgRP) and tyrosine hydroxylase (TH) markers. Collectively, these findings indicate that the non-AgRP/TH Arc^GABA^ PVH^CRH^ neurocircuit serves as a critical neural substrate that directly encodes the scalability of stress responses to environmental stressors by modulating inhibitory GABA release in a stimulus intensity-dependent manner.

## 1. Introduction

Stress exposure inevitably disrupts normal physiological homeostasis across species^1^, and an appropriate stress response is essential for responding to, coping with, and subsequently recovering from those disturbing stimuli^2,3^. Dysregulated responses to environmental stressors often result in abnormal psychiatric and behavioral outcomes and, when severe, can lead to stress-related mental illnesses such as anxiety disorders and posttraumatic stress disorder (PTSD)^1,4,5^. Currently, due to the lack of adequate treatments, stress-related mental disorders represent a significant public health concern. Therefore, to develop effective therapies for the prevention and treatment of these disorders, it is critically important to fully understand how the brain precisely encodes the appropriate scalability of stress responses.

Stress responses are mediated by the hypothalamic-pituitary-adrenal (HPA) axis, whose activity is governed by corticotropin-releasing hormone (CRH) neurons in the paraventricular nucleus of the hypothalamus (PVH, denoted as PVH^CRH^ neurons)^2,6,7^. Upon exposure to stress stimuli, PVH^CRH^ neurons are rapidly activated, resulting in the prompt release of CRH into the pituitary portal circulation and subsequent activation of the HPA axis ^8–10^. Notably, the level of PVH^CRH^-HPA axis activity plays a pivotal role in determining the magnitude of the stress response. Accordingly, transient activation of PVH^CRH^ neurons is sufficient to evoke an acute stress response in unstressed animals, whereas selective silencing of these neurons rapidly attenuates stress responses in stressed animals^10,11^. PVH^CRH^ neuronal activity is tightly regulated by a balance between excitatory and inhibitory actions imposed by upstream GABAergic and glutamatergic inputs, as well as other hormonal and neuromodulatory afferents^12–14^. It has been estimated that more than one-third of the synaptic inputs onto PVH^CRH^ neurons are GABAergic^15^, underscoring the critical role of inhibitory circuits in regulating PVH^CRH^-mediated stress responses. In contrast to the excitatory effects of glutamatergic inputs, GABAergic inputs play a dominant role in restraining PVH^CRH^ neuronal responses to stress cues^7,8,12,15–17^, highlighting their potential as therapeutic targets for stress-related mental disorders, which are often associated with hightened PVH^CRH^ neuronal activity. Although the dynamic recruitment of PVH^CRH^ neurons and their downstream outputs during stress has been well characterized^7,8,17,18^, relatively little is known about how upstream GABAergic inputs directly gate PVH^CRH^ neurons to encode and process stress-related information.

Emerging evidence indicates that stress-related emotional states are closely intertwined with feeding regulation by hypothalamic neurons, suggesting that shared neurocircuits co-regulate stress and feeding^19–25^. Neurons in the arcuate nucleus (Arc), a well-established feeding-driving center, provide dense GABAergic inputs to PVH neurons^26–30^. Despite extensive evidence demonstrating the regulation of Arc GABAergic neurons (denoted as Arc^GABA^ neurons) and Arc^GABA^→PVH projections in feeding behaviors^28–34^, the direct gating of stress responses by PVH-projecting Arc^GABA^ neurons, particularly via direct Arc^GABA^→PVH^CRH^ projections, has not been previously investigated.

In the present study, we first established, using pseudorabies viral retrograde tracing, that Arc^GABA^ neurons represent a major GABAergic upstream input to PVH^CRH^ neurons. We then found that, although PVH^CRH^ neurons exhibited time-locked activation in response to various environmental stressors, both GABA release onto PVH^CRH^ neurons and PVH^CRH^-projecting Arc^GABA^ neuronal activity were reduced under prolonged, high-intensity stress conditions, but not following acute, low-intensity stress exposure. Intriguingly, selective manipulation of PVH^CRH^-projecting Arc^GABA^ neuronal activity resulted in corresponding changes in both PVH^CRH^-mediated stress responses and HPA axis outputs, phenocopying the effects of direct manipulation of PVH^CRH^ neuronal activity. Furthermore, we observed that these PVH^CRH^-projecting Arc^GABA^ neurons are directly regulate feeding behaviors, while not belonging to AgRP- or tyrosine hydroxylase (TH)-expressing subsets. Together, our data reveal a novel neurocircuit in which non-AgRP/TH Arc^GABA^ neurons gate inhibitory GABA release onto PVH^CRH^ neurons in a stimulus intensity-dependent manner, thereby regulating stress scalability. These findings identify a promising candidate neural substrate for managing stress-related psychiatric disorders.

## 2. Results

### 2.1 PVH^CRH^ neurons receive monosynaptic functional GABAergic inputs from Arc^GABA^ neurons

PVH^CRH^ neurons are known to receive inputs from multiple upstream targets, including inhibitory and excitatory projections, as well as other hormonal and neuromodulatory tones^12–14^. To better understand how upstream neurons form monosynaptic inputs onto PVH^CRH^ neurons, we delivered a mixture of AAVDJ-CAG-FLEX-TVA-mCherry and AAVDJ-EF1A-DIO-oG-WPRE-hGH into the PVH of *Crh-ires-cre* mice, followed by a second injection of pseudorabies viruses EnvA RV-GFP into the same location four weeks later. GFP-positive neurons were identified in many brain regions (**Fig.1a-d**), including the ventral portion of the bed nucleus of the stria terminalis (vBNST), the dorsomedial hypothalamic nucleus (DMH), the lateral section of the parabrachial nucleus (LPBN), the median preoptic area (mPOA) and the supraoptic nucleus (SON). Notably, among these regions, the Arc area, a well-established hypothalamic GABAergic center that sends inhibitory inputs to the PVH, contained the most robust number of retrogradely labelled neurons compared with other identified GABAergic regions (*e.g.,* DMH, mPOA, and vBNST^12,35,36^).

**Fig. 1.**
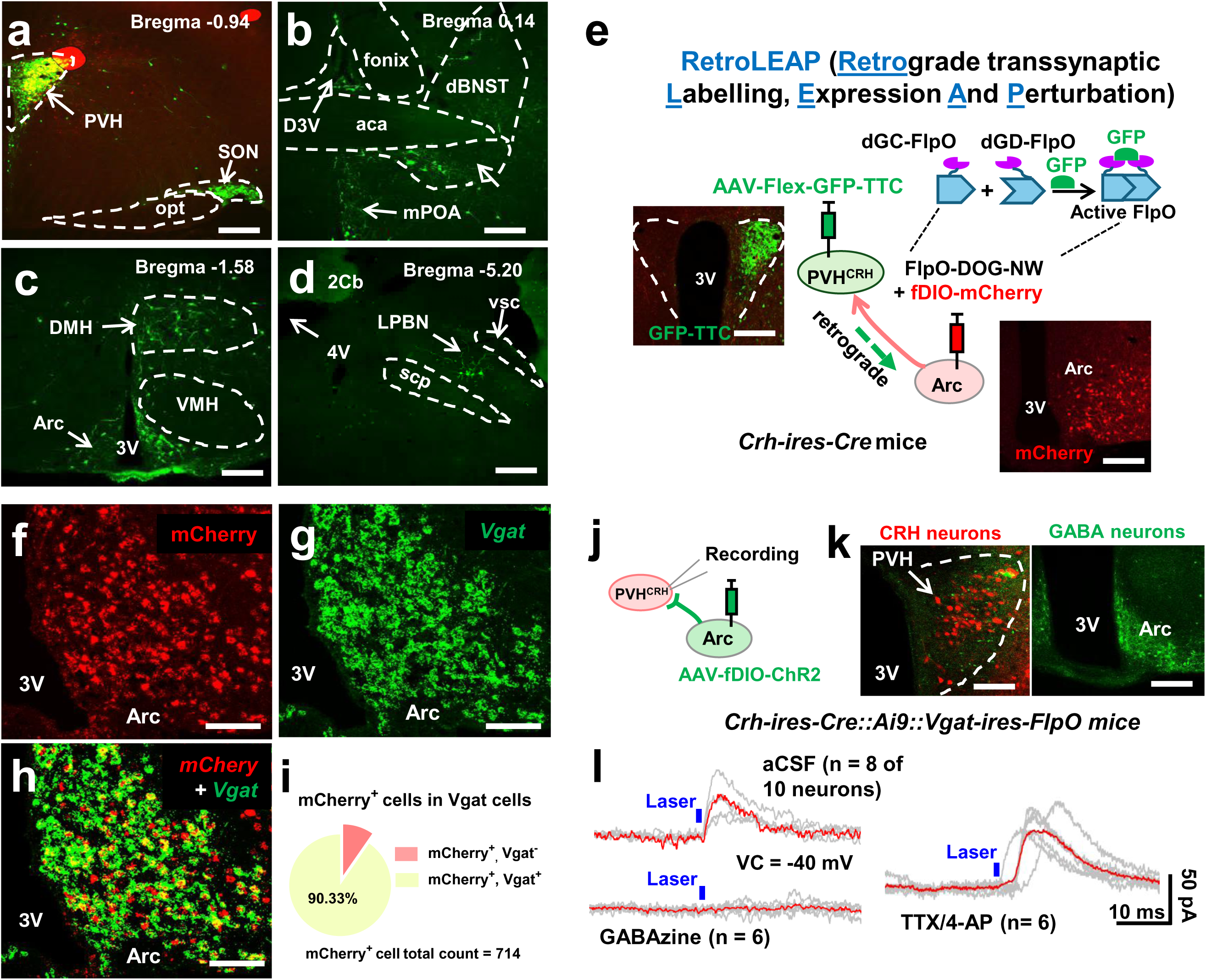
ArcGABA neurons represent a major source of inhibitory input to PVH^CRH^ neurons. (**a-d**) Psudotyped rabies tracing shows the key upstream targets of PVH^CRH^ neurons. (**e**) Experimental schematic showing the working design of the RetroLEAP approach, and representative images showing GFP-TTC (green) expression in the PVH and mCherry (red) expression in the Arc. (**f-i**) Representative images of RNAScope ISH showing that majority of Arc neurons retrogradedly labelled from PVH^CRH^ neurons (*mCherry*, red) express *Vgat* (green) , and quantificative analyses confirmed around 90% of *mCherry*-expressing *cells* are also *Vgat* positive (total 714 cells in 3 mice). (**j**) Experimental schematic of whole cell voltage-clamp recording. (**k**) Representative images showing ChR2 expression (green) in Arc^GABA^ cells, and CRH reporter neurons (red) innervated by ChR2-expressing fibers. (**l**) Example whole cell voltage-clamp recording traces showing oIPSCs evoked by 1-ms light pulses in 8 of 10 recorded reporter PVH^CRH^ neurons. All evoked oIPSCs were blocked by application of GABA receptor antagonist GABAzine, whereas refractory to treatment of polysynaptic blockers 4-AP/TTX (n = 3 mice). aca: anterior commersiure; Arc: arcuate nuclus; dBNST: dorsal bed nucleus of the stria terminalis; DMH: dorsamedial nucleus of hypothalamus; D3V: dorsal 3rd ventricle; 2cb: second cerebellum lobule; 3V: the 3rd ventricle; 4V: the 4th ventricle; LPBN: lateral paeabrachial nucleus; mPOA: medial preoptic area; PVH: paraventricular nucleus of hypothalamus; scp: superior cerebellar peduncle; VMH: ventromedial nucleus of hypothalamus; vsc: ventral spinocerebellar tract. Scale bars: 250 µm in **a-d**; 100 µm in **e** and right image of **k**; 20 µm in **f-h** and left image of **k**.

To confirm that Arc^GABA^ neurons form monosynaptic inputs onto PVH^CRH^ neurons, we employed the RetroLEAP (<u>Retro</u>grade transsynaptic <u>L</u>abeling, <u>E</u>xpression <u>A</u>nd <u>P</u>erturbation) technique (**Fig.1e**), which is designed to trace presynaptically connected upstream neurons by injecting a non-toxic, Cre-dependent retrograde GFP-TTC (non-toxic heavy-chain of the tetanus toxin) vector into Cre-expressing starter downstream neurons, followed by delivery of a GFP-dependent flip-recombinase (FlpO-DOG^37^) together with FlpO-dependent reporter or effector vectors^38–40^. Briefly, in *Crh-ires-Cre* mice, AAV2/DJ8-CAG-DIO-EGFP-TTC was injected into the PVH, followed by injecting a mixture of AAV-EF1a-FlpO-DOG-NW and FlpO-dependent AAV-Ef1a-fDIO-mCherry into the Arc. Six weeks later, double fluorescent *in situ* hybridization (ISH) was performed using RNAscope probes against *mCherry* and *Vgat*. As shown in **Fig.1f-d**, *Vgat* expression colocalized with the vast majority of the PVH^CRH^-retrogradely labelled, *mCherry*-expressing Arc neurons, demonstrating a predominantly GABAergic connection between these neuronal populations.

To further confirm the functional connection between PVH^CRH^ and Arc^GABA^ neurons, we first recorded inhibitory postsynaptic currents (IPSCs) in PVH^CRH^ neurons following photostimulation Arc^GABA^ fibers. To achieve this, a channelrhodopsin (ChR2)-eGFP fusion protein (AAV-fDIO-ChR2-eGFP) was expressed in the Arc of *Cre-ires-Cre::Rosa-lsl-tdTomato::Vgat-FlpO* mice (**Fig.1j, k**). 8 out of 10 recorded PVH^CRH^ neurons exhibited robust light-induced postsynaptic responses. These IPSCs were completely abolished by bath application of the GABA-A receptor blocker GABAzine (n = 6), but were refractory to the treatment of 4-AP/TTX, indicating the monosynaptic nature of the connetion between Arc^GABA^ neurons and PVH^CRH^ neurons (**Fig.1l**). We next measured GABA release onto PVH^CRH^ neurons in response to specific activation of their upstream Arc^GABA^ neurons. To accomplish this, we expressed a Cre-dependent AAV-hSyn-Flex-iGABASnFR sensor and retrograde GFP-TTC vectors in the PVH, followed by expression of AAV-FlpO-DOG together with a FlpO-dependent AAV-nEF-Coff/Fon-ChRmine-oScarlet vector in the Arc of *Cre-ires-Cre* mice (**Supplementary Fig.1a**). *In vivo* fiber photometry recording revealed a significant increase in iGABASnFR signals in PVH^CRH^ neurons during photostimulation (635 nm wave length) of ChRmine-expressing Arc^GABA^ fibers (**Supplementary Fig.1b**), indicating a time-locked increase in GABA release onto PVH^CRH^ neurons following Arc^GABA^ neuron activation. Taken together, these results demonstrate that Arc^GABA^ neurons provide a major source of monosynaptic GABAergic inputs to PVH^CRH^ neurons.

### 2.2. GABA release and neuronal activity within the Arc^GABA^ PVH^CRH^ neurocircuit scale with environmental stressor intensity

Previous studies have demonstrated that PVH^CRH^ neurons exhibit robust, time-locked increase in activity in response to stress exposure, and that their activity states directly reflect the expression of stress-related behaviors^10,11,21^. Given that our tracing results identified Arc^GABA^ neurons as a major source of GABAergic input to PVH^CRH^ neurons, these findings suggest that the Arc^GABA^→PVH^CRH^ projection may play a key role in directly regulating stress responses. To assess the contribution of this neurocircuit to the sensing and processing of stress-related signals, we monitored *in vivo* GABA release and neuronal activity within this circuit in freely behaving mice (**Fig. 2a**). To this end, using the RetroLEAP approach, we virally expressed Cre-dependent AAV-hSyn-Flex-iGABASnFR sensor and retrograde GFP-TTC vectors in the PVH, followed by expression of AAV-EF1a-FlpO-DOG-NW together with FlpO-dependent AAV-Ef1a-fDIO-GCaMP6s or AAV-Ef1a-fDIO-sRGECO vectors in the Arc of *Crh-ires-Cre* mice. Simultaneous *in vivo* fiber photometry recordings were then performed using a branching bundle patch cord, with one fiber-optic cannula implanted above the PVH and the other implanted above the Arc with a 10° angle, in response to multiple aversive stressors, including loud sound, air puff, water spray, 15 seconds (s) tail-restraint test (TRT), and 30 s physical restraint^10,21,22,39,41^. An additional group of *Crh-ires cre* mice with GCaMP6m calcium indicator expression and fiber-optic canula implantation only in the PVH was also included as a positive control to record stress-induced PVH^CRH^ neuronal activity.

**Fig. 2.**
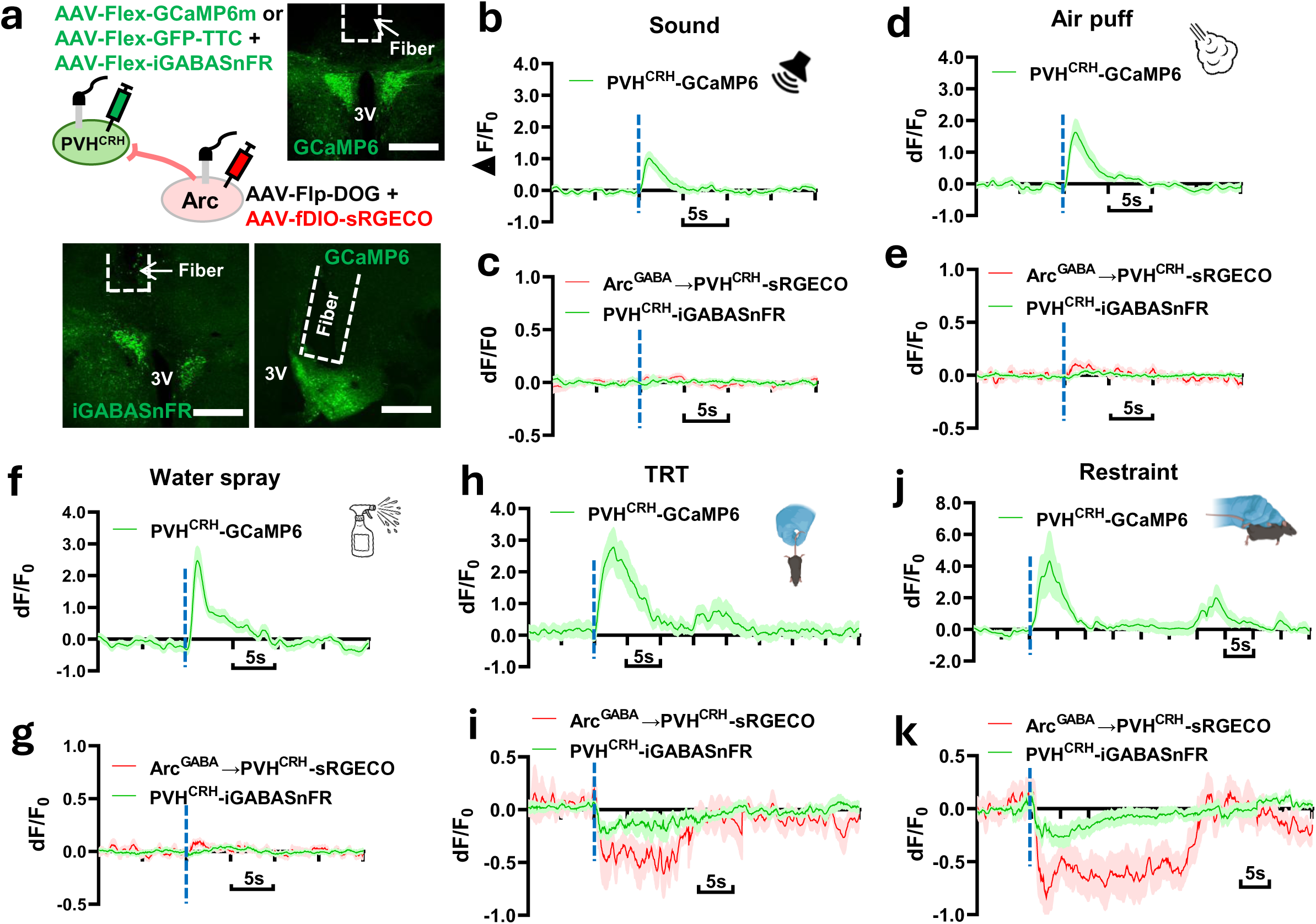
GABA transmission and neuronal activity within the Arc^GABA^ PVH^CRH^ circuit scale with environmental stressor strength. (**a**) Experimental schematic and representative images showing GCaMP6 and iGABASnFR (green) expression in the PVH, GCaMP6 (green) expression in the Arc, and optic fiber implantations above the PVH and Arc. Histograms of Ca^2+^ signals in PVH^CRH^ neurons (**b, d, f, h, j**) and concurrent iGABA signals in PVH^CRH^ neurons and Ca^2+^ signals in PVH^CRH^-projecting Arc^GABA^ cells (**c, e, g, i, k**), aligned to the onset of stressors: lound sound (**b** and **c**), air puff (**d** and **e**), water spray (**f** and **g**), 15 s TRT (**h** and **i**), and 30 s restraint (**j** and **k**). 3V: the 3rd ventricle. Scale bars: 250 µm in **a**.

Our fiber photometry recordings showed that, consistent with previous results^10,21^, PVH^CRH^ neuronal GCaMP6 signals exhibited robust, time-locked increase in freely behaving mice in response to all tested aversive stimuli (**Fig. 2b,d,f,h,j**). In contrast to this stressor-dependent activation of PVH^CRH^ neurons, no obvious changes were detected in either iGABASnFR signals or GCaMP6 signals in PVH^CRH^-projecting Arc^GABA^ neuronal in response to aversive stimuli triggered by loud sound, air puff, or water spray (**Fig. 2c,e,g**). However, both signals showed a robust, time-locked reduction following tail suspension and restraint stress (**Fig. 2i,k**), which were associated with longer-lasting (*e.g.,* Δ*F/F* increase duration) and stronger (*e.g.,* Δ*F/F* peak value) activation of PVH^CRH^ neurons compared with those induced by loud sound, air puff, or water spray. These observations demonstrate that changes in GABA release onto PVH^CRH^ neurons, as well as activity dynamics in PVH^CRH^-projecting Arc^GABA^ neurons, are selectively recruited by stress signals that elicited prolonged, high-intensity responses. Importantly, this response strength-dependent manner of recruitment indicates that neurotransmission and neuronal activity within the Arc^GABA^→PVH^CRH^ neurocircuit scale with environmental stressor intensity.

### 2.3. PVH^CRH^-projecting Arc^GABA^ neurons exhibit a stressor-dependent activity anticorrelation with PVH^CRH^ neurons

While it was unexpected that GABA release onto PVH^CRH^ neurons, along with PVH^CRH^-projecting Arc^GABA^ neuronal activation, exhibited a selective response pattern to different stressors, it remains unclear whether this non-ubiquitous recruitment of the Arc^GABA^→PVH^CRH^ projection depends on the perceived scalability or intensity of the stress signals. To directly address this question, we simultaneously measured activity dynamics of both PVH^CRH^-projecting Arc^GABA^ neurons and PVH^CRH^ neurons in response to the same aversive stressors. To this end, using the RetroLEAP approach, we virally expressed Cre-dependent GCaMP6m or jRGEco1a calcium indicator and retrograde GFP-TTC vectors in the PVH, followed by expression of AAV-EF1a-FlpO-DOG-NW in combination with FlpO-dependent AAV-Ef1a-fDIO-GCaMP6s vectors in the Arc of *Crh-ires-Cre* mice (**Fig. 3a**). Simultaneous *in vivo* fiber photometry recordings were then performed, as described above, during exposure to multiple aversive stressors.

**Fig. 3.**
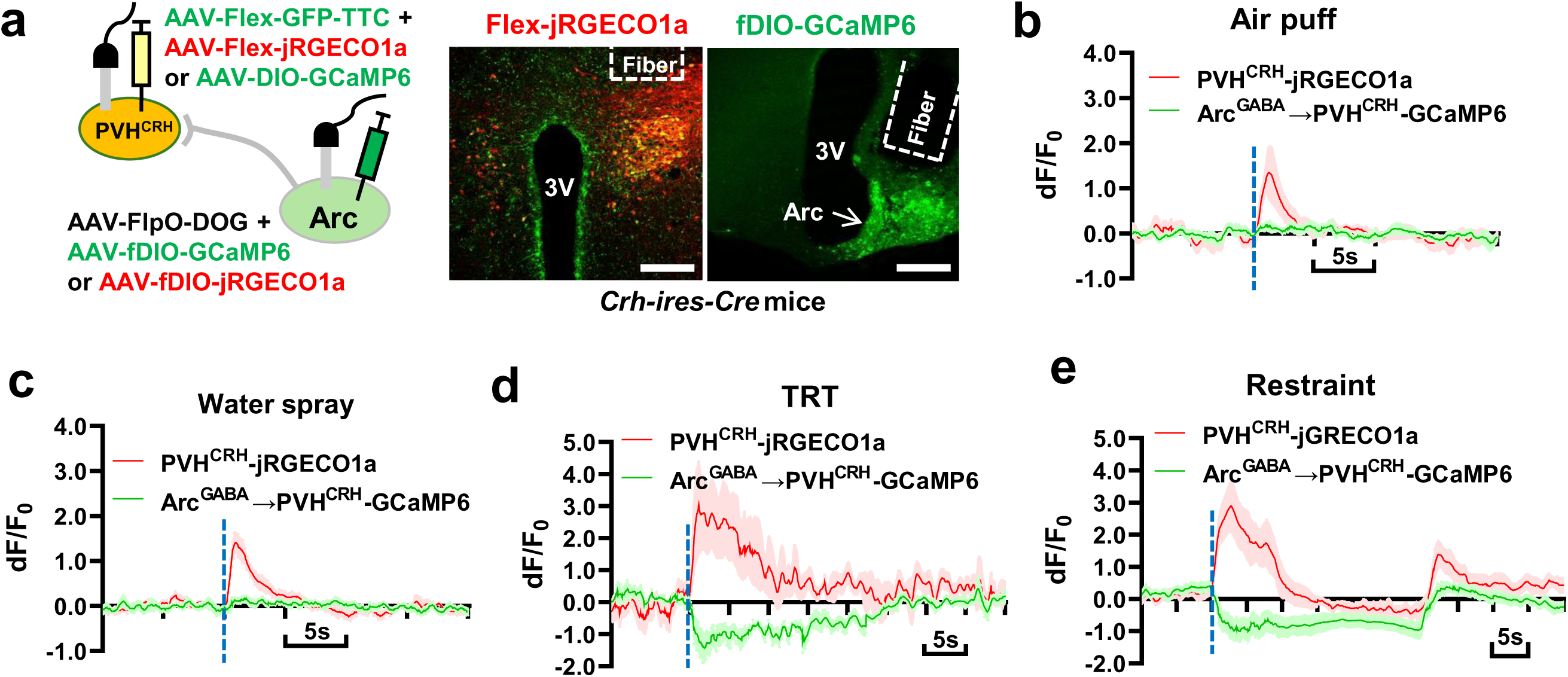
Activity dynamics between PVH^CRH^ and PVH^CRH^-projecting Arc^GABA^ neurons are correlated in response to the same stressor. (**a**) Experimental schematic and representative images showing Cre-dependent GFP-TTC and jGRECO1a (yellow) expression in the PVH, and Flp-dependnet GCaMP6 (green) expression in the Arc, and optic fiber implantations above the PVH and Arc. (**b-e**) Histograms of concurrent Ca^2+^ signals in PVH^CRH^ neurons (red traces) and PVH^CRH^-projecting Arc^GABA^ cells (green traces), aligned to the onset of stressors: air puff (**b,** n = 6), water speay (**c,** n = 6), 15 s TRT (**d,** n = 8), and 30 s restraint (**e,** n = 12). 3V: the 3rd ventricle. Scale bars: 250 µm in **a**.

Whereas time-locked increases were ubiquitously observed in PVH^CRH^ neurons in response to all tested stress stimuli, no obvious changes were detected in concurrently recorded PVH^CRH^-projecting Arc^GABA^ neurons following exposure to loud sound, air puff, or water spray(**Supplemeanty Fig. 2**; **Fig. 3b** & **c**). In contrast, in response to the same TRT and restraint stress, PVH^CRH^-projecting Arc^GABA^ neuronal activity exhibited a robust, time-locked anticorrelation with PVH^CRH^ neuronal activity, paralleling the reduction in GABA release onto PVH^CRH^ neurons (**Fig. 3c** & **d**). These observations demonstrate that selective anticorrelated changes in PVH^CRH^-projecting Arc^GABA^ neuronal activity play a critical role in orchestrating behavioral responses to prolonged, high-intensity stressors. Collectively, this stressor-dependent coordination of neuronal activity highlights the existence of a precise encoding mechanism within the Arc^GABA^→PVH^CRH^ neurocircuit that underlies appropriate stress responses.

### 2.4. Activity states of PVH^CRH^-projecting Arc^GABA^ neurons directly influence HPA axis output, stress levels and anxiety-like behaviors

GABAergic transmission-mediated regulation (e.g., inhibition or activation) of the HPA axis at the level of PVH^CRH^ neurons has been well established. To determine whether changes in the activity of PVH^CRH^-projecting Arc^GABA^ neurons directly mirror those of PVH^CRH^ neurons, we selectively activated or inhibited this neuronal population using a chemogenetic approach and examined the resulting effects on stress levels and HPA axis output.

Firstly, we tested the direct effects of silencing PVH^CRH^-projecting Arc^GABA^ neurons by virally expressing GCaMP6 calcium indicator in the PVH and inhibitory Designer Receptors Exclusively Activated by Designer Drugs (DREADDs) in the Arc of *Crh-ires-cre* mice using the RetroLEAP strategy (**Fig. 4a**). Five weeks later, under non-stress conditions, blood samples were collected following intraperitoneal (i.p.) administration of either saline or deschloroclozapine (DCZ,1 µg/Kg body weight^42^), with one week apart between treatments, and plasma Cort levels were measured. As shown in **Fig. 4b**, DCZ treatment resulted in a significant increase in plasma Cort levels compared with saline administration. We next examined whether simultaneous inhibition of PVH^CRH^-projecting Arc^GABA^ neurons, which reduces GABA release, enhances stress-induced activation of PVH^CRH^ neurons in response to aversive stimuli. Mice were conditioned to the same stressors, and PVH^CRH^ neuronal activity was recorded under either saline or DCZ treatment. As shown in **Fig. 4c-j**, stressor-induced increases in PVH^CRH^ neuronal activity were significantly augmented under DCZ treatment compared with saline controls. We then tested whether selective silencing of PVH^CRH^-projecting Arc^GABA^ neurons produces anxiogenic effects. Compared with control mice, chemogenetic inhibition of PVH^CRH^-projecting Arc^GABA^ neurons resulted in a significant reduction in time spent in both the open arms of the elevated plus maze (EPM) test and in the light compartment of the light/dark (LD) box (**Fig. 4k & l**). Finally, c-Fos immunostaining confirmed that inhibition of PVH^CRH^-projecting Arc^GABA^ neurons led to robust activation of PVH^CRH^ neurons (**Fig. 4m**). Together, these functional studies demonstrate that a reduction in intrinsic activity of PVH^CRH^-projecting Arc^GABA^ neurons is sufficient to induce a stress-prone state, likely mediated through a fast-acting, HPA-dependent pathway.

**Fig. 4.**
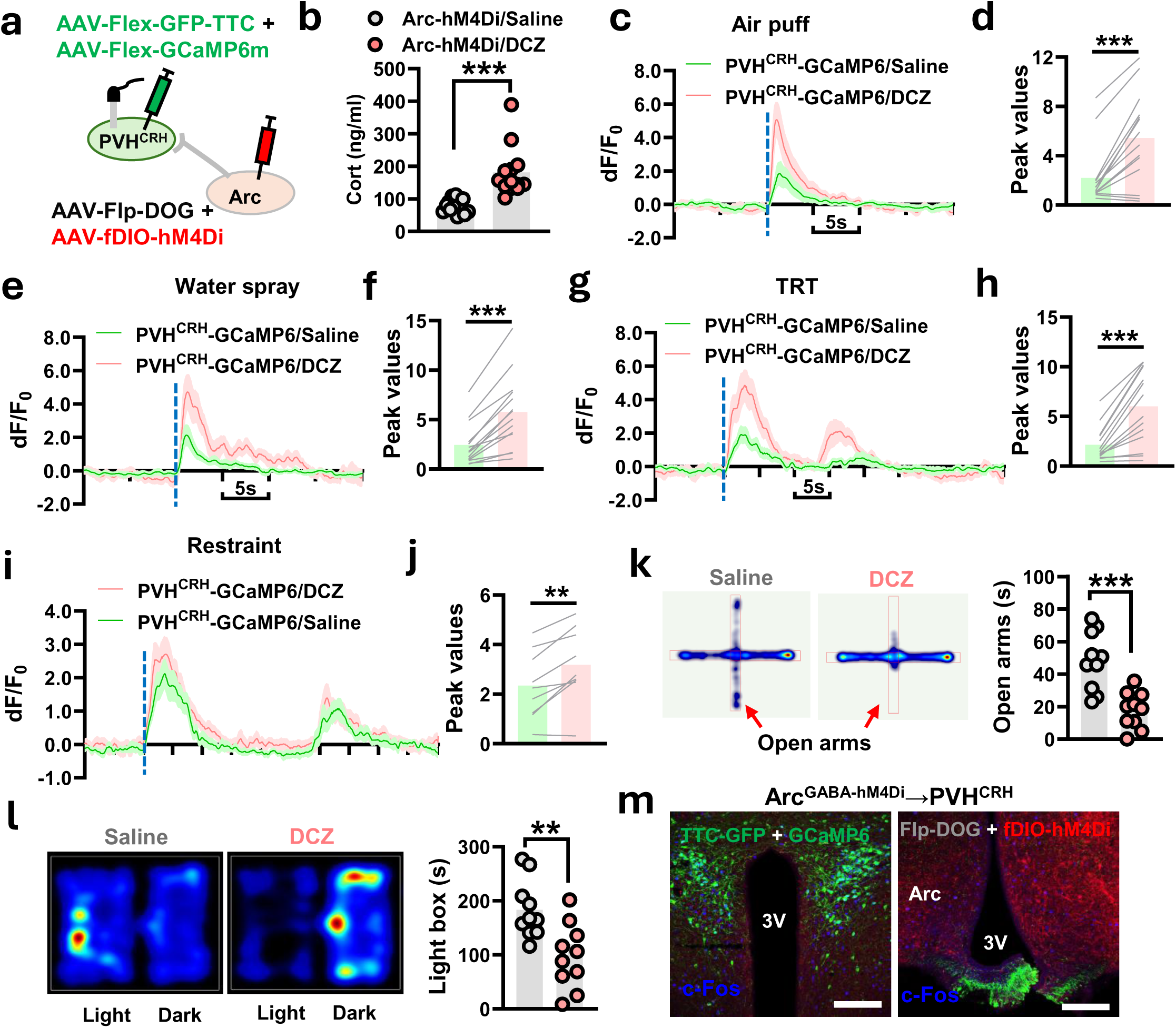
Silencing of PVH^CRH^-projecting Arc^GABA^ neurons augments stress responses and activates HPA axis output. (**a**) Experimental schematic. (**b**) Plasma Cort levels at 1 h after either saline (gray) or DCZ (red) injection (n = 13*** *p* < 0.001, unpaired *t*-test, all values are presented as mean ± S.E.M.). (**c-j**) Histograms and peak value comparisons of Ca^2+^ signals in PVH^CRH^ neurons under saline (green traces) or DCZ (red traces) treatment condition, aligned to the onset of stressors: air puff (**c & d,** n = 14; *** *p* < 0.001, paired *t*-test), water spray (**e & f,** n = 14; *** *p* < 0.001, paired *t*-test), 15 s TRT (**g & h,** n = 14; *** *p* < 0.001, paired *t*-test), and 30 s restraint (**i & j,** n = 9; ** *p* < 0.01, paired *t*-test). (**k**) Representive trace of EPM test and quantification of time spent in open arms (n = 10; *** *p* < 0.001, paired *t*-test, all values are presented as mean ± S.E.M.). (**l**) Representive trace of LD box test and quantification of time spent in light compartment (n = 10; *** *p* < 0.001, paired *t*-test, all values are presented as mean ± S.E.M.). (**m**) Representative images showing the expression GFP-TTC/GCaMP6 (green), AAV-fDIO-hM4Gi (red), and c-Fos (blue) in the PVH and Arc area in the PVH and Arc area at 1 h post-DCZ treatment . Arc: arcuate nucleus; 3V: the 3rd ventricle. Scale bars: 100 µm in **m**.

We next tested the direct effects of activating PVH^CRH^-projecting Arc^GABA^ neurons by virally expressing the GCaMP6 calcium indicator in the PVH and excitatory DREADDs in the Arc of *Crh-ires-cre* mice using the RetroLEAP approach (**Fig. 5a**). Compared with non-stress conditions, plasma Cort levels were robustly elevated by 30-min restraint stress in saline-treated mice; however, this stress-induced increase was nearly completely abolished by DCZ treatment applied 15 min prior to restraint stress (**Fig. 5b**). Consistent with these endocrine changes, chemogenetic activation of PVH^CRH^-projecting Arc^GABA^ neurons significantly suppressed stressor-induced increases in PVH^CRH^ neuronal activity and produced in an anxiolytic effect, as evidenced by the significantly increased time spent in both the open arms of the EPM test and the light compartment of the LD box test (**Fig. 5c-l**). Collectively, these results demonstrate that increased intrinsic activity of PVH^CRH^-projecting Arc^GABA^ neurons is sufficient to induce a stress-resistant state, likely through suppression of HPA axis activity.

**Fig. 5.**
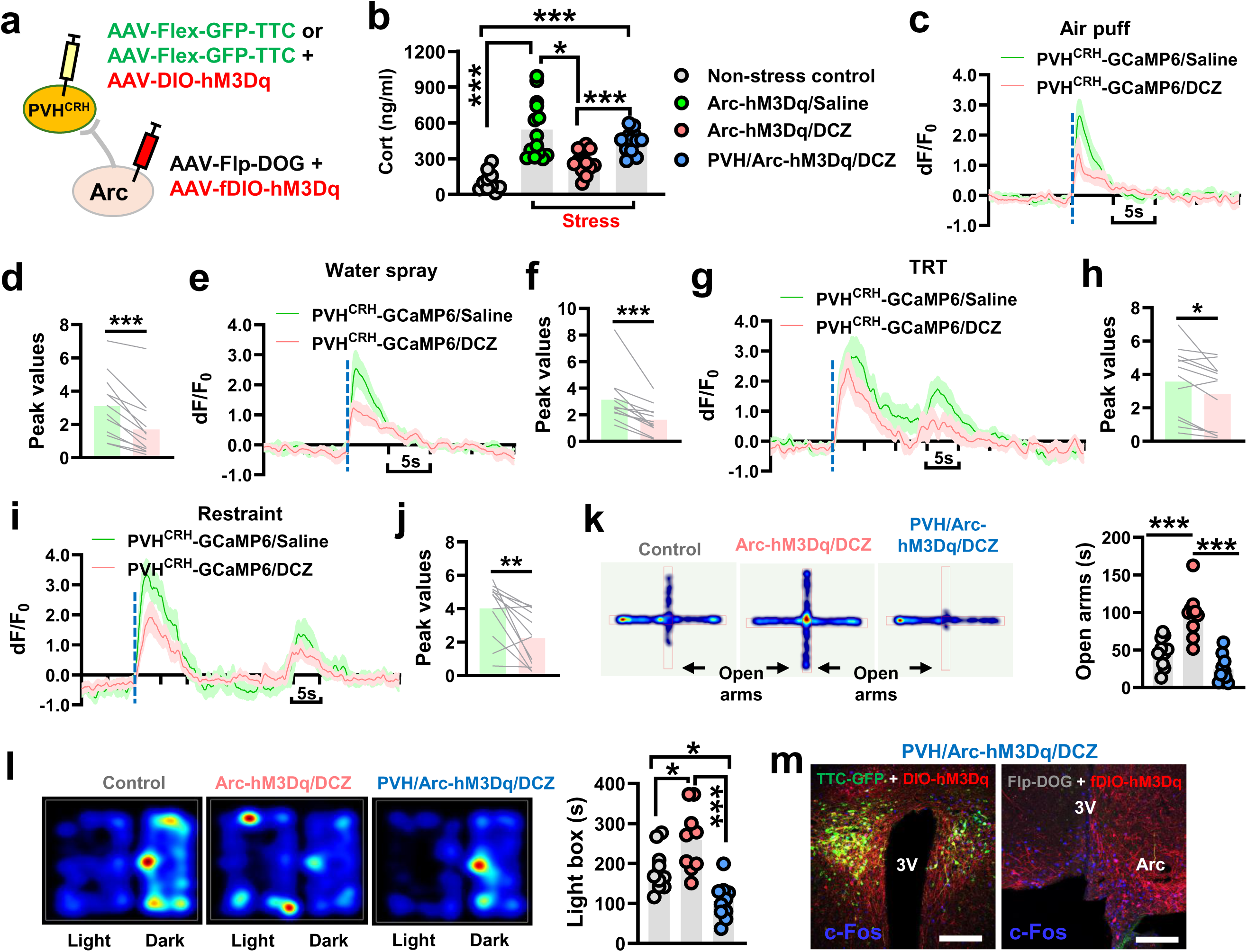
Activation of PVH^CRH^-projecting Arc^GABA^ neurons attenuates stress responses and suppresses HPA axis output. (**a**) Experimental schematic. (**b**) Plasma Cort levels under non-stress control condition or at 30 min following 30-min restraint stress under either saline (gray) or DCZ (red) treatment (n = 10 – 14; * *p* < 0.05; *** *p* < 0.001, Two-way ANOVA, all values are presented as mean ± S.E.M.). (**c-j**) Histograms and peak value comparisons of Ca^2+^ signals in PVH^CRH^ neurons under saline (green traces) or DCZ (red traces) treatment condition, aligned to the onset of stressors: air puff (**c & d,** n = 12; *** *p* < 0.001, unpaired *t*-test, all values are presented as mean ± S.E.M.), water spray (**e & f,** n = 12; *** *p* < 0.001, paired *t*-test), 15 s TRT (**g & h,** n = 12; * *p* < 0.05, paired *t*-test), and 30 s restraint (**i & j,** n = 12; ** *p* < 0.01, paired *t*-test). (**k**) Representive trace of EPM test and quantification of time spent in open arms (n = 10; *** *p* < 0.001, One-way ANOVA, all values are presented as mean ± S.E.M.). (**l**) Representive trace of LD box test and quantification of time spent in light compartment (n = 10; * *p* < 0.05; *** *p* < 0.001, One-way ANOVA, all values are presented as mean ± S.E.M.). (**m**) Representative images showing the expression of GFP-TTC/ hM3Gq (yellow), AAV-fDIO-hM3Gq (red), and c-Fos (blue) in the PVH and Arc area at 1 h post-DCZ treatment. Arc: arcuate nucleus; 3V: the 3rd ventricle. Scale bars: 100 µm in **m**.

Given that Arc^GABA^ neurons, including those expressing Agouti-related peptide (AgRP), send collateral projections to multiple downstream targets^31,43^, it is possible that other redounant neurocircuits may play a more predominant role in mediating the observed regulation of stress responses by PVH^CRH^-projecting Arc^GABA^ neurons. To further test the direct invovlement of the Arc^GABA^→PVH^CRH^ projection in gating stress levels and HPA axis output, we examined whether the effects of increased activity in PVH^CRH^-projecting Arc^GABA^ neurons, which promotes GABA release, are occuluded by concomitant activation of PVH^CRH^ neurons. To this end, we virally expressed excitatory DREADDs in both the PVH and Arc of *Crh-ires-Cre* mice (**Fig. 5a**). As shown in **Fig. 5b**, the suppressing effects of selective activation of PVH^CRH^-projecting Arc^GABA^ neurons on endocrine changes, as well as its anxiolytic effects, were diminished by simultaneous excitation of PVH^CRH^ neurons (**Fig. 5k** & **l**), indicating that PVH^CRH^ neurons directly mediate the downstream effects of Arc^GABA^ activation. Notably, concurrent excitation of PVH^CRH^ neurons exerted a dominant functional effect, as mice exhibited elevated plasma Cort levels and increased anxiety compared with controls (**Fig. 5b, k, l**). These findings, together with the partial reversal of stressor-triggered PVH^CRH^ neuronal activation by selective activation of PVH^CRH^-projecting Arc^GABA^ neurons, suggest that additional inhibitory upstream inputs beyond Arc^GABA^ neurons are required to fully constrain PVH^CRH^-mediated stress responses. Finally, c-Fos immunostaining confirmed effective chemogenetic activation in both targeted neuronal populations across all tested mice (**Fig. 5m**).

### 2.5. PVH^CRH^-projecting Arc^GABA^ neurons represent an unique subset of Arc neruons also involve in feeding regulation

Arc^GABA^ neurons are known to comprise diverse cell subpopulations, including agouti-related protein (AgRP)/Neuropeptide Y neurons, dopaminergic neurons, proopiomelanocortin (POMC) neurons, non-AgRP/non-POMC neurons expressing Cre driven by the rat insulin-2 promoter (RIP-Cre) or the pancreas duodenum homeobox 1 promoter (Pdx1-Cre), and neurons expressiong kisspeptin, growth hormone-releasing hormone or dopamine, among others^27,31,43–46^. Many of these neuronal populations are know to play important roles in feeding regulation. We then tested whether PVH^CRH^-projecting Arc^GABA^ neurons also regulate feeding behavior. To this end, we measured food intake in mice following chemogenetic activation of PVH^CRH^-projecting Arc^GABA^ neurons during light cycle, when mice are typically in a sated, calorically replete state. Compared with the control condition, mice with PVH^CRH^-projecting Arc^GABA^ neuronal activation consumed a significantly higher amount of food during the 4 h test window (**Fig. 6a-c**). These data demonstrate that PVH^CRH^-projecting Arc^GABA^ neurons directly participate in the regulation of feeding behavior.

**Fig. 6.**
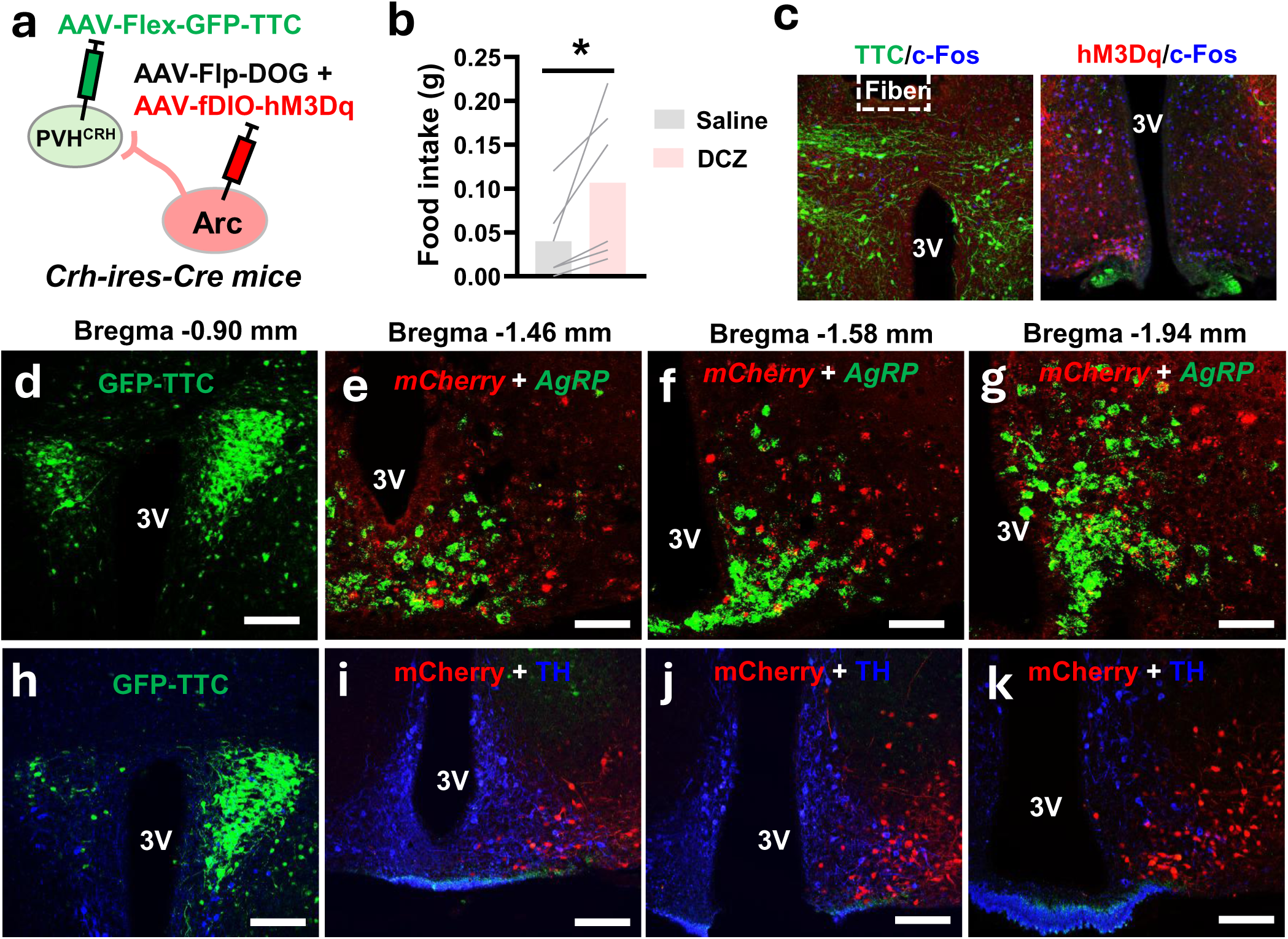
PVHCRH-projecting Arc^GABA^ neurons are non-AgRP/TH feeding-regulatory subet. (**a**) Experimental schematic. (**b**) Food intake during the light cycle (9AM -1PM, n = 6; * *p* < 0.05; paired *t*-test). (**c**) Representative images showing the c-Fos expression (blue) in the PVH and Arc area at 1 h post-DCZ treatment. (**d -g**) Representative images showing GFP-TTC expression in the PVH, and *mCherry* (red) and *AgRP* (green) mRNA expression in the Arc of male *Crh-ires-cre* mice (**h -k**) Representative images showing GFP-TTC (green), *mCherry* (red), and TH (blue) expression in the PVH and Arc area of male *Crh-ires-cre* mice. Scale bars: 100 µm in **c, d** and **h-k**, 20 µm in **e-g**.

To better understand which subtype(s) of Arc^GABA^ neurons provide the direct GABAergic input to PVH^CRH^ neurons responsible for gating the scalibity of stress responses, we then employed RetroLEAP approach combined with RNAScope ISH and immunofluorescent stainings to define the molecular profiles of PVH^CRH^-projecting Arc^GABA^ neurons. Our RNAscope ISH analysis further revealed that vast majority of PVH^CRH^-projecting Arc neurons were negative for *AgRP* expression, while only a small proportion of them co-express *neurotensin (Nts)* and *Pomc* (**Fig. 6d-g**, Supple. **Fig. 2**). Furthermore, we found that PVH^CRH^-retrogradely labeled Arc neurons did not express tyrosine hydroxylase (TH), a marker for dopaminergic neurons (**Fig. 6h-k**).These findings suggest that PVH^CRH^-projecting Arc neurons belong to a non-AgRP/TH neuronal subset.

## 3. Discussion

In this study, using RetroLEAP neurocircuit targeting approaches, chemogenetics, and fiber photometry, we revealed a novel hypothalamic inhibitory circuit that encodes the scalability of stress responses. Specifically, we demonstrate that Arc^GABA^ neurons send monosynaptic GABAergic inputs to PVH^CRH^ neurons, modulating GABAergic transmission onto PVH^CRH^ neurons and altering their activity dynamics in a stressor duration- and strength-dependent manner during stress exposure (**Fig. 7**). In addition, we found that the intrinsic activity of PVH^CRH^-projecting Arc^GABA^ neurons directly gauges HPA axis output, associated axiety states, and feeding drive (**Fig. 7**). Anatomically, these neurons represent a non-AgRP/TH neuronal subset. Together, these data suggest that the Arc^GABA^→PVH^CRH^ circuit serves as an important neural substrate for achieving allostatic stress states.

**Fig. 7.**
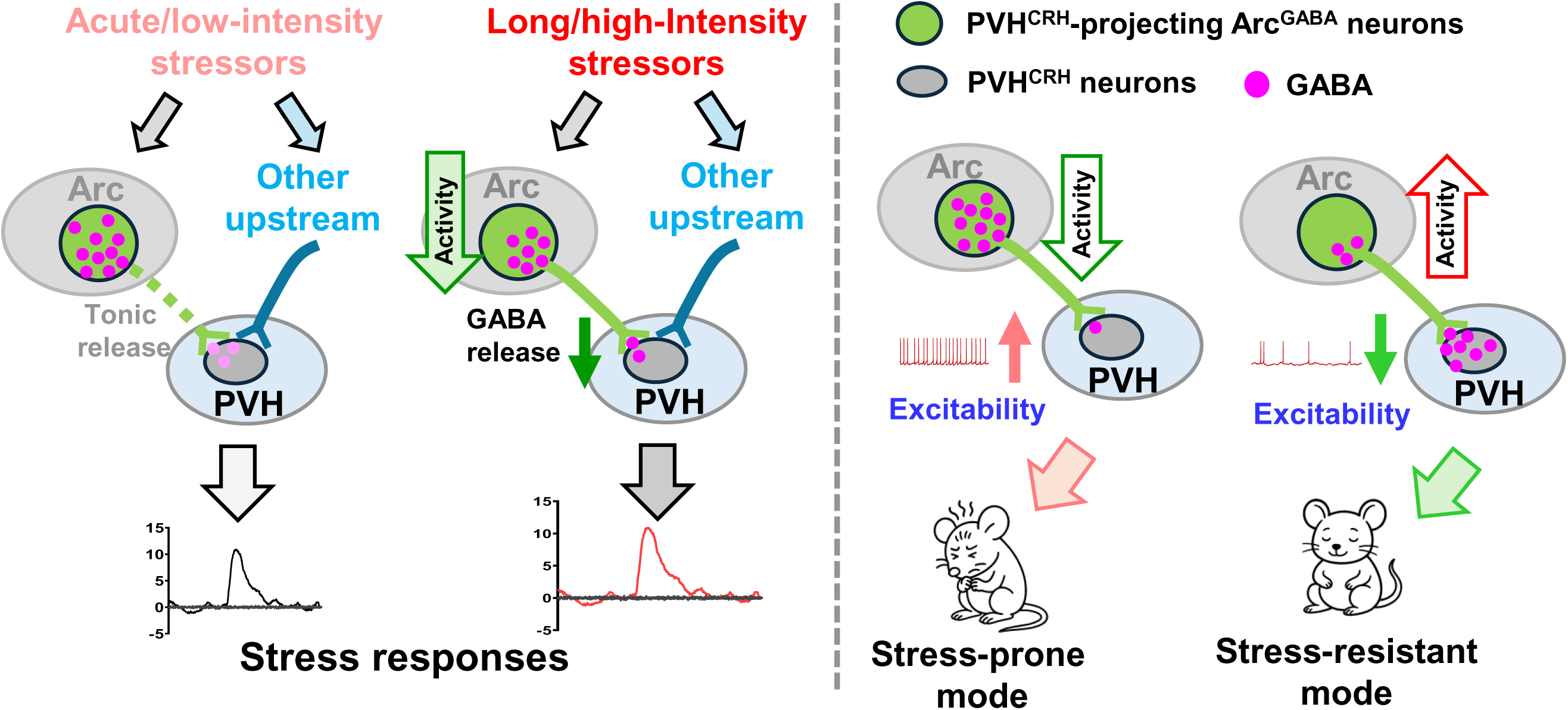
Model summarizing how dynamic Arc^GABA^→PVH^CRH^ circuit activity encodes the appropriate scalibility of stress-related responses. Under basal condtions, PVH^CRH^-projecting Arc^GABA^ neurons provide tonic GABA inputs to PVH^CRH^ neurons and are not specifically recruited during exposure to transient, low-intensity stressors (e.g., air puff). In contrast, these neurons are recuited during prolonged, high-intensity stressors (i.e., 30 s restraint), leading to a time-locked reduction in neuronal activity and GABA release onto downstream PVH^CRH^ neurons, thereby triggering an appropriate stress response. Phenocoping the effects of direct manipulation of PVH^CRH^ neuronal activity, intrinsic activity levels of PVH^CRH^-projecting Arc^GABA^ neurons directly gate HPA axis output and stress-related emotional states.

It is well-established that GABAergic inputs play a dominant role in restraining PVH^CRH^ neuronal activity in response to stress cues^7,8,12,15–17^. Consistent with this, local administration of the GABA_A_ receptor antagonist increases plasma Cort levels as well as CRH gene transcription and c-Fos expression in the PVH^47,48^, whereas microinfusion of GABA agonists, such as the stress-derived neurosteroid, into the PVH decreases circulating Cort levels^49^. Furthermore, the number of GABAergic synapses terminating on PVH^CRH^-positive profiles increases by approximately 55% following adrenalectomy, a manipulation that elevates the synthetic and secretory activity of CRH neurons^15^. Notably, GABAergic innervation to PVH^CRH^ neurons are traditionally considered to originate primarily from neurocircuits located adjacent to the PVH, including the peri-PVH zone, neighboring forebrain regions, as well as that within the PVH^16,47,50^. This raises the question of whether non-adjacent PVH^CRH^-projecting Arc^GABA^ neurons also play a direct role in regulating PVH^CRH^-mediated stress responses, as previous studies suggest that they may instead act indirectly via relays through these local GABAergic circuits^36,51^. Consistent with prior evidence that Arc neurons provide dense GABAergic inputs to the PVH^26–30^, our current results demonstrate that Arc^GABA^ neurons serve as a major functional source of GABAergic input to PVH^CRH^ neurons. Importantly, this Arc^GABA^→PVH^CRH^ projection directly impacts stress responses by modulating GABA release and neuronal activity dynamics in a stressor-dependent manner. In addition, in line with the well-established role of Arc^GABA^ neurons on feeding regulation^33,52^, we found that this PVH^CRH^-projecting Arc^GABA^ neuronal population is also regulating feeding behavior. Together, these findings highlight the Arc^GABA^→PVH^CRH^ circuit as a shared neural pathway that co-regulates stress and feeding^19–25^.

While the dynamic recruitment of PVH^CRH^ neurons and their outputs during stress has been well characterized^7,8,17,18^, how PVH^CRH^-projecting inhibitory inputs dynamically contribute to stress-related signal processing remains largely unknown. In particular, it is unclear how each source of these inputs directly orchestrates PVH^CRH^-mediated neuroendocrine responses to achieve appropritate scalability of stress expression. It has been suggested that activation of CRH neurons occurs alongside a reduction in inhibitory GABAergic transmission, thereby lowering the threshold for neuronal firing upon exposure to stress cues^16,36^. While we observed activation of PVH^CRH^ neurons in response to all tested stressors, our experiments detected a pronounced decrease in GABA release onto PVH^CRH^ neurons only in response to aversive stressors of prolonged duration and higher intensity (i.e., 15 s TRT, or 30 s restraint). Given that *in vivo* fiber photometry is a highly sensitive optical technique used for recording real-time neural activity^53^, the absence of a detectable reduction in GABA release onto PVH^CRH^ neurons in response to transient stressors (i.e., loud sound, air puff, or water spray) in this study may reflect that these stimuli are either insufficiently intense or not of appropriate modality to reach the threshold required to alter GABA transmission, including this directly contributed by Arc^GABA^ neurons. Importantly, these findings provide novel evidence indicating that the reduction of inhibitory GABAergic transmission in CRH neurons is not an uniform event but instead depends on the nature of the stressor. In a pattern correlated with reduced GABA release onto PVH^CRH^ neurons, the time-locked decrease in the activity of PVH^CRH^-projecting Arc^GABA^ neurons in response to TRT and restraint stress suggests that Arc^GABA^-derived inhibitory input is specifically recruited to gate response strength to aversive stressors of prolonged duration and higher intensity. Notably, this specific recruitment pattern highlights a coordinated role for the PVH^CRH^-projecting inhibitory network in regulating the scalability of stress responses.

While hormonal regulation has long been extensively studied, recent findings highlight the critical role of synaptic inputs from upstream neurons in modulating PVH^CRH^-mediated stress responses^11,54^, underscoring the importance of top-down neuromodulation in shaping stress-related emotional adaptations. Chronic stress, a major driver of mental disorders, is often associated with impaired GABAergic inhibition on PVH^CRH^ neurons^15,16,48^. However, the origin of this inhibitory deficient remains largely unknown. Additionally, the direct impacts of activity changes in PVH^CRH^-projecting GABAergic upstreams on HPA axis outputs and emotional states have yet to be fully determined. Consistent with their role in dynamically orchestrating allostatic stress responses, our findings show that the activity sates of PVH^CRH^-projecting Arc^GABA^ neurons bidirectionally influence circulating Cort levels and anxiety-like behaviors, demonstrating their essential role in maintaining balanced HPA axis function. Therefore, these results indicate that the Arc^GABA^→PVH^CRH^ circuit represents a critical top-down neuromodulatory mechanism that gates appropriate responposes to envirobmental stressors, and that its dysfunction may direclty contributes to the development of stress-related mental disorders.

In line with emerging evidence that stress responses and feeding regulation share common neural substrates^19–25^, a notable and concerning aspect of many psychotropic agents used to treat anxiety and depression is their strong tendency to induce hunger, weight gain, and metabolic disease^55^. Our findings are particularly important becasue chemogenetic activation of PVH^CRH^-projecting Arc^GABA^ neurons promotes feeding behavior, a change that is associated with a stress- and anxiety-resistant state. This suggests that the Arc^GABA^→PVH^CRH^ neurocircuit may contribute to the development of the “obese” phenotype observed during anti-psychiatic treatment. From a pharmacologic perspective, the mechanistic demonstration of a direct link between orexingeric Arc^GABA^ neurons and anxiogenic PVH^CRH^ neurons provides a foundational framework for understanding this previously difficult-to-define behavioral phenomenon in humans. Especially, our results further indicate that the Arc^GABA^→PVH^CRH^ neurocircuit could serve as a promising therapeutic target for developing effective and safe strategies to mitigate treatment-associated weight gain during the management of stress-related mental disorders.

The complexity of Arc^GABA^ neurons and their extensive communication with neurons both within and ouside the Arc have been well established^44^. Despite their orexigenic role, most PVH^CRH^-projecting Arc^GABA^ neurons express neither AgRP nor TH, markers of two key Arc neuronal populations known to promote feeding^33,46^. This finding is consistent with previous studies showing that AgRP neurons, although they send inhibitory projections to the PVH and attenuate behavioral indices of stress^31,32,34^, do not provide direct input to PVH^CRH^ neurons^29,30,56^. Notably, we also confirmed that only a small subset of PVH^CRH^-projecting Arc^GABA^ neurons are either POMC or Nts positive. While the Arc^POMC^→PVH^CRH^ has been implicated in mediating HPA axis activation during either physical restraint or negative energy balance^57,58^, our experimental design did not allow us to isolate the specific contribution of these POMC^+^ neurons, as we broadly targeted all PVH^CRH^-projecting Arc neurons. Similarly, despite its known anorexigenic role^59^, the exact function of Nts^+^ PVH^CRH^-projecting Arc^GABA^ neurons in orchestrating stress-related behaviors remains to be clearly defined. Additionlly, the application of scRNA-Seq will be essential to determine whether these PVH^CRH^-projecting Arc expressing Arc^GABA^ neurons express other Arc markers, such as the leptin receptor^60^. A clear defination of the molecular identity of this neuronal population will open new avenues for targeting this circuit in the development of effective therapeutics for stress-related mental disorders.

In summary, our findings demonstrate that a subset of non-AgRP/TH Arc^GABA^ neurons, serving as a major source of GABAergic input, are directly recuited to transmit environmental stress-related information to PVH^CRH^ neurons in a stimulus strenth-dependent manner. This inhibitory Arc^GABA^→PVH^CRH^ neurocircuit represents a key substrate for dynamically encoding the scalibity of stress responses, and its functional deficit may directly contribute to the development of stress-related mental disorders.

## 4. Materials and Methods

### Animals

Mice were housed at 21°C -22°C with a 12 h light/12 h dark cycle with standard pellet chow and water provided *ad libitum*. Animal care and procedures were approved by the University of Texas Health Science Center at Houston Institutional Animal Care and Use Committee. *Crh-ires*_-_*Cre* (JAX:012704), *Ai9* reporter (JAX:007909), and *Vgat-FlpO* mice (JAX:031331) mice were purchased from the Jax lab and described and confirmed to express Cre in PVH^CRH^ neurons previously^61,62^. Both male and female mice were used as study subjects, except otherwise noted. All mice that were used for stereotaxic injections were 8-10 weeks old. All study subjects were littermates and randomly distributed between study groups. Group size was estimated based on relevant literature reports and differences in average between groups from pilot studies.

### Surgeries and viral constructs

Briefly, mice were anesthetized with a ketamine/xylazine cocktail (100 mg/kg and 10 mg/kg, respectively), and their heads affixed to a stereotaxic apparatus. Viral vectors were delivered through a 0.5 µL syringe (Neuros Model 7000.5 KH; Hamilton, Reno, NV, USA) mounted on a motorized stereotaxic injector (Quintessential Stereotaxic Injector; Stoelting, Wood Dale, IL, USA) at a rate of 40 nL/min. Viral preparations were titered at ∼10^12^ particles/mL. Viral delivery was targeted to the PVH (anteroposterior (AP): -0.90 mm; mediolateral (ML): ± 0.10 mm; dorsoventral (DV): -4.90 mm) or Arc (AP: -1.40 mm and -1.60 mm; ML: ± 0.20 mm; DV: -5.90 mm).

For mapping the upstream neurons that make monosynaptic inputs onto PVH^CRH^ neurons, a mixture of AAVDJ-CAG-FLEX-TVA-mCherry and AAVDJ-EF1A-DIO-oG-WPRE-hGH (Salk Institute) were delivered into the bilateral PVH (200 nl/side) of *Crh-ires-cre* mice. Four weeks later, a second injection of pseudo rabies viruses EnVA RV-GFP was delivered to the same location (Salk Institute, 800 nl/side). Mice were perfused 10 days later.

For measuring the GABA release in PVH^CRH^ neurons in response to the activation of their upstream Arc^GABA^ neurons, we employed the RetroLEAP (retrograde transsynaptic Labeling, Expression And Perturbation) technique in *Crh-ires-Cre* mice. This technique is designed to trace presynaptically connected upstream neurons by injecting the non-toxic Cre-dependent retrograde GFP-TTC (non-toxic heavy-chain of the tetanus toxin) vector into Cre-expressing starter downstream neurons, and followed by injecting a mixture of flip-recombinase whose activity depends on the presence and binding of GFP (FlpO-DOG^37^), along with FlpO-dependent interested vectors^38–40^. Briefly, AAV2/DJ8-CAG-DIO-EGFP-TTC (Canadian Neurophotonics Platform, SCR_016477) together with AAV-hSyn-Flex-iGABASnFR.F102G (Addgene, plasmid # 112164) was injected bilaterally into the PVH (200 nl/each side), followed by injecting a mixture of AAV-EF1a-FlpO-DOG-NW (Addgene, plasmid # 75469) and FlpO-dependent AAV-nEF-Coff/Fon-ChRmine-oScarlet (Addgene, plasmid # 137160) vectors into the bilateral Arc (1:1, 200 nl/each side; AP: -1.50 mm; ML: ±0.20 mm; DV: -5.90 mm) of *Crh-ires-cre* mice. Then, we implanted the fiber-optic cannula (Ø1.25 mm, Ø400 µm Core, 0.39 NA; Doric Lens) above the above the PVH.

For recording the activity changes of PVH^CRH^ neurons, AAV-Syn-Flex-GCaMP6m-WPRE.SV40 (Addgene, plasmid # 100838) was injected bilaterally into the PVH (150 nl/each side) of *Crh-ires-cre* mice, followed by implanting a single fiber optic cannula (Ø1.25 mm Stainless Ferrule, Ø400 µm Core, 0.39 NA; Doric Lens, Québec, Canada) above the PVH (AP: -0.90 mm; ML: ±0.0 mm; DV: -4.80 mm).

For simultaneously recording the GABA release onto PVH^CRH^ neurons and the activity changes of PVH^CRH^-projecting Arc^GABA^ neurons, AAV2/DJ8-CAG-DIO-EGFP-TTC together with AAV-hSyn-Flex-iGABASnFR.F102G was injected bilaterally into the PVH (200 nl/each side), followed by injecting a mixture of AAV-EF1a-FlpO-DOG-NW and FlpO-dependent AAV-Ef1a-fDIO-GCaMP6s (Addgene, plasmid # 105714) vectors into the bilateral Arc (1:1, 200 nl/each side) of *Crh-ires-cre* mice. Then, we implanted the first fiber-optic cannula above the above the PVH, and the second one above the Arc area with a 10° angle (AP: -1.50 mm; ML: -1.30 mm; DV: -5.80 mm).

For simultaneously recording the activity dynamics of PVH^CRH^ neurons and PVH^CRH^-projecting Arc^GABA^ neurons, a mixture of AAV2/DJ8-CAG-DIO-EGFP-TTC and AAV-Syn-Flex-GCaMP6m-WPRE.SV40 was bilaterally injected into the PVH (1:1, 200 nl/each side), followed by bilateral injection of a mixture of AAV-EF1a-FlpO-DOG-NW and FlpO-dependent AAV-Ef1a-fDIO-GCaMP6s vectors into the Arc (1:1, 200 nl/each side) of *Crh-ires-cre* mice. Subsequently, the first fiber-optic cannula was implanted above the PVH, and the second canula was placed above the Arc with a 10° angle as described above.

For testing the direct impacts of changing activity states of PVH^CRH^-projecting Arc^GABA^ neurons on stress levels and HPA axis output, a combination of AAV2/DJ8-CAG-DIO-EGFP-TTC and AAV-Syn-Flex-GCaMP6m-WPRE.SV40 (1:1, 200 nl/each side) was injected into the bilateral PVH (150 nl/each side), followed by injecting a mixture of AAV-EF1a-FlpO-DOG-NW and FlpO-dependent AAV-Ef1a-fDIO-hM3Dq-mCherry or AAV-Ef1a-fDIO-hM4Di-mCherry (Baylor Viral Core) into the bilateral Arc (1:1, 200 nl/each side) of *Crh-ires-cre* mice.

For testing whether the direct role of PVH^CRH^-projecting Arc^GABA^ neurons on controlling stress levels and HPA axis output is mediated via PVH^CRH^ neurons, a mixture of AAV2/DJ8-CAG-DIO-EGFP-TTC and AAV-hSyn-DIO-hM3D(Gq)-mCherry (Addgene, plasmid # 44361) was injected into the bilateral PVH (200 nl/each side), and followed by injecting a mixture of AAV-EF1a-FlpO-DOG-NW and FlpO-dependent AAV-Ef1a-fDIO-hM3D(Gq)-mCherry into the bilateral Arc (1:1, 200 nl/each side) of *Crh-ires-cre* mice.

### Brain slice electrophysiological recordings

Coronal brain slices (250–300 μm) containing the PVH from mice that had received stereotaxic injections of AAV-fDIO-ChR2-eGFP to Arc (4 weeks after injection) were cut in ice-cold artificial cerebrospinal fluid (aCSF) containing the following (in mm): 125 NaCl, 2.5 KCl, 1 MgCl_2_, 2 CaCl_2_, 1.25 NaH_2_PO_4_, 25 NaHCO_3_, and 11 D-glucose bubbling with 95% O_2_/5% CO_2_. Slices containing the LS were immediately transferred to a holding chamber and submerged in oxygenated aCSF. Slices were maintained for recovery for at least 1 h at 32–34 °C before transferring to a recording chamber. Individual slices were transferred to a recording chamber mounted on an upright microscope (Olympus BX51WI) and continuously superfused (2 ml/min) with ACSF warmed to 32-34 °C by passing it through a feedback-controlled in-line heater (TC-324B; Warner Instruments). Cells were visualized through a × 40 water-immersion objective with differential interference contrast optics and infrared illumination. Whole cell voltage-clamp recordings were made from neurons within the LSv that showed the highest density of ChR2-eYFP^+^ axonal fibers. Patch pipettes (3–5 MΩ) were filled with a Cs^+^-based low Cl^−^ internal solution containing (in mm) 135 CsMeSO_3_, 10 HEPES, 1 EGTA, 3.3 QX-314, 4 Mg-ATP, 0.3 Na_2_-GTP, 8 Na_2_-Phosphocreatine (pH 7.3 adjusted with CsOH; 295 mOsm) for voltage–clamp. For current–clamp recordings, pipettes were filled with a K^+^-based low Cl^−^ internal solution containing (in mm) 145 KGlu, 10 HEPES, 0.2 EGTA, 1 MgCl_2_,4 Mg-ATP, 0.3 Na_2_-GTP, 10 Na_2_-Phosphocreatine (pH 7.3 adjusted with KOH; 295 mOsm). Membrane potentials were corrected for ∼10 mV liquid junction potential. To activate ChR2 light from a 473 nm laser (Opto Engine LLC, Midvale, UT, USA) was focused on the area of the recorded PVH^CRH^ reporter neurons to produce spot illumination through optic fiber. Brief pulses of light (473 nm blue light, 1-2 mW/mm^2^) were delivered at the recording site at 15 s or 200 ms intervals under control of the acquisition software. GABAzine (10 µm) drugs (Abcam) were bath-applied to block GABA-A receptors during voltage-clamp recordings of photostimulation-induced inhibitory current responses. TTX (0.5 μm; Alomone labs, Jerusalem, Israel), and 4-AP (100 μm; ACROS Organics, Fisher Scientific, Pittsburgh, PA, USA) were bath-applied during voltage-clamp recordings of photostimulation-induced inhibitory current responses to block action potentials and inhibit network activity.

### Fiber photometry

*In vivo* recordings were performed using the R821/FR-21Tricolor Multichannel Fiber Photometry System (RWD life science, Shenzhen, China) in an open-top home cage. All tests were conducted during the light cycle following a minimum 4 weeks recovery period post-surgery, and the animal ID was blinded when recordings were conducted. Briefly, 560-, 470- and 410-nm laser beams were first launched into the fluorescence cube and then coupled into the optical fibers; the 410-nm laser served as a motion-control reference. For simultaneous dual sites recordings, a branching bundle patch cord with 2 outputs, one connected with fiber canula implanted above the PVH and the other once connected with fiber canula implanted above the Arc, were used (Doric Lens, Québec, Canada). GCaMP6/iGABASnFR signals and control emission fluorescence were collected by the camera at the sample rate of 20 Hz.

For recording GABA release onto in PVH^CRH^ neurons in response to photostimulation of Arc^GABA^ inputs, animals were habituated in the recording cage for 10 minutes (min) with the optic patch fiber connected. After habituation, iGABASnFR signals were recorded for 1 min. At 30 s after the start of recording, photostimulation was delivered for 10 s using a 635 nm laser source of red light (RWD life science) at a frequency of 5Hz and a power intensity of 20mW.

For the air puff stimulus, a brief buff of compressed air (20 psi) was delivered via a nozzle positioned ∼3 cm from the mouse, and recording continued for ∼ 2 mins. For water spray stimulus, a brief stimulus was delivered by spraying water once toward the head of the mouse using a sprayer, and recording continued for ∼ 2 mins. For tail restraint tests, mice were suspended in the air for 15 s by gently grabbing the tail and then released back into their home cages, and recording lasted for ∼ 2 mins. For physical restraint stress, mice were gently grabbed and restrained for 30 s with hands and then released back into their home cages, and recording continued for ∼ 2 mins. Different stimuli were delivered at intervals of at least 10 min.

For single-cannula recordings in the PVH or the Arc, animals were habituated in the recording cage for 10 min with the optic patch fiber connected. After habituation, signals and mice behaviors were recorded for 10 min. The first stimulus was delivered 2 min after the start of recording, and subsequent stimuli were presented at intervals of at least 10 min.

For dual-cannula recordings in the PVH and the Arc, animals were habituated in the recording cage for 10 min with the optic fiber connected. After habituation, signals and mice behaviors were recorded for 10 min. The first stimulus was delivered 2 min after the start of recording, and subsequent stimuli were presented at intervals of at least 10 min. The air puff, water spray, tail suspension, and restraint stress were presented to mice on different dates, at least three days apart.

For single-cannula recordings in the PVH combined with chemogenetic manipulations of PVH^CRH^-projecting Arc^GABA^ neuronal activities in the Arc, animals were injected with either saline or DCZ 20 min before habituation, and were then habituated in the recording cage for 10 min with the optic patch cable connected. After habituation, signals and mice behaviors were recorded for 10 min. The first stimulus was delivered 2 min after the start of recording, and subsequent stimuli were presented at intervals of at least 10 min.

The photometry signal (*F*) was calculated as the ratio *F_470_/F_410_*, and ΔF/F was computed as (F – F_0_)/F_0_, where F_0_ represents the median value of the photometry signal. Only calcium signals >3 standard deviations were considered events. For each mouse, ΔF/F values were averaged across multiple trials for subsequent analysis.

### Behavioral assay

All behavioral assays were performed after a 4-week recovery period from surgery, and animal identities were blinded during behavior tests. All mice were tested using the open field exploration test (OFT), elevated plus-maze (EPM) test and light-dark (LD) box test, which are well-established paradigms for assessing stress-associated anxiety-like behaviors in rodents^63–67^.

### OFT

The OFT apparatus consists of a white Plexiglas box (size: 40 × 40 × 40 cm) placced in a low-noise testing room with stable lighting conditions. After a 30-min acclimation, each mouse was placed in a corner of the arena to initiate a 20_-_min test session. Between trials, the apparatus was thoroughly cleaned with 70% ethanol and allowed to dry completely.

### EPM

The EPM (Kinder Scientific, Poway, CA, USA), consisting of two open and two closed arms, was positioned 50 cm above the floor in a low-noise testing room with stable lighting. After a 30-min acclimation, each mouse was placed on the central platform facing an open arm, to initiate a 10_-_min session test. The maze was thoroughly cleaned with 70% ethanol and dried between trials.

### LD box test

The LD box test apparatus consists of two compartments (Kinder Scientific, Poway, CA, USA), with the light chamber occupying two-thirds of the total area and the dark chamber occupying the remaining one-third. After 30 min acclimation, each mouse was placed in the dark room with free access to the light compartment and allowed to explore freely for 10 min. The apparatus was thoroughly cleaned with 70% ethanol and dried between trials.

All tests were simultaneously video recorded (Noldus, Leesburg, VA, USA), and behavioral indices were measured with an automated video tracking system (Ethovision XT). Time spent in the center of OFT, in the open arms of the EPM, and in the light compartment of the LD box was quantified and compared between experimental groups.

### Blood sample collection and measurement

In all mice, blood samples were collected between 9:00 AM and 12:00 PM using the submandibular bleeding approach. In mice that received hM4D(Gi) viral delivery in the Arc, blood samples were collected 1 h following the application of either saline or DCZ under non-stress conditions. In mice that received hM3D(Gq) viral delivery in the Arc, blood samples were collected 30 min following restraint stress and 1 h after administration of either saline or DCZ. In mice that received hM3D(Gq) viral delivery in both the Arc and the PVH, blood samples were collected 30 min following restraint stress and 1 h after administration of either saline or DCZ. All blood samples were then measured for corticosterone (Cort) levels, the direct readout of HPA axis activity, by the Vanderbilt University Medical Center (VUMC) Analytical Services Core, as described in our previous work^21^. Blood samples collected from mice that received neither viral injection nor restraint stress were also measured for Cort levels and served as normal controls.

### RNAscope ISH

For in situ hybridization experiments, as described in our previous studies^39,68^, fixed-frozen brain tissues were sectioned at 15 μm thickness using a sliding microtome and mounted onto Superfrost Plus Gold slides (Fisher Scientific, INC). Sections were first air dry at room temperature (RT) for 30 minutes and baked at 60 °C for 30 minutes. Sections were post-fixed with pre-cold 10% neutral buffered formalin at 4 °C for 15 minutes. Sections were further dehydrated with 50%, 70%, and 100% ethanol at RT and then applied with 3% hydrogen peroxide for 10 minutes at RT. After 10 minutes’ antigen retrieval at 98_∼_100 °C, sections were digested with protease (RNAscope™ Protease III, #322340, Advanced Cell Diagnostic, INC). Sections were hybridized with probes against *Vgat* (RNAscope® Probe-Mm-Slc32a1, # 319191-C3, Advanced Cell Diagnostic, INC), *mCherry* (RNAscope® Probe-Mm-mCherry-O2, # 409021-C2, Advanced Cell Diagnostic, INC), *Agrp* (RNAscope® Probe-Mm-Agrp, # 400711-C1, Advanced Cell Diagnostic, INC), *Pomc* (RNAscope® Probe-Mm-Pomc, # 519151-C1, Advanced Cell Diagnostic, INC), and *Neurotensin* (RNAscope® Probe-Mm-Nts, # 420441-C3, Advanced Cell Diagnostic, INC) for 2 h at 40 °C and signals were amplified with RNAscope® Multiplex Fluorescent Reagent Kit v2 with TSA Vivid Dyes (#323270, Advanced Cell Diagnostic, INC).

### Immunohistochemistry (IHC)

After all behavioral tests, mice were anesthetized with ketamine/xylazine (150 mg/kg and 15 mg/kg, respectively). After loss of the pedal reflex, mice were transcardially perfused with 15 ml of saline and 15 ml of 10% buffered formalin (In Vivo Perfusion System IV-140, Braintree Scientific Inc.). The brains were then collected and stored in 10% buffered formalin overnight at room temperature, and then, the brain was switched to 30% sucrose in PBS for overnight. Brains were sectioned into 30 μM coronal slices on a frozen sliding microtome and stored in 0.1% NaN3 in PBS at 4 °C.

For IHC, sectioned slices were rinsed with 0.3% Triton X-100 in phosphate-buffered saline (PBS) for 5 min three times and were blocked in 0.3% Triton X-100 in PBS with 10% donkey serum at room temperature (RT) for 1 hour. The slices were incubated in a primary antibody solution (primary antibody, 5% donkey serum, and 0.3% Triton X-100 in PBS) overnight at 4°C.

The following primary antibodies were used: c-Fos rabbit monoclonal antibody (mAb) (9F6) (1:1000, #2250; Cell Signaling Technology), tyrosine hydroxylase rabbit polyclonal antibody (1:1000, ab112; Abcam). For secondary antibody treatment, slices were rinsed with 0.3% Triton X-100 in PBS for 5 min for three times and incubated in the secondary antibody solution (Alexa Fluor 647–conjugated AffiniPure Donkey (H+L) anti-rabbit immunoglobulin G (Jackson ImmunoResearch), 1:400, 10% donkey serum, and 0.3% Triton X-100 in PBS) for 2 hours at RT. The floating slices were mounted onto microscope slides and coverslipped with Fluoromount (Diagnostic BioSystems Inc., Sigma-Aldrich). A confocal microscope was used to image the slices at different resolutions (Leica TCS SP5, Leica Microsystems, Wetzlar, Germany). Mice with offsite injections and cannula implants were removed from the study.

### Quantification and Statistic analyses

GraphPad Prism 10.2.3 (GraphPad Software, Inc., La Jolla, CA, USA) was used for all statistical analyses and construction parts of the figures. For fiber photometry data, raw data from single trials was first processed through pMat application (https://github.com/djamesbarker/pMAT).86 Individual traces were aligned to zero at stimulus onset and averaged to be visualized in MATLAB R2022b (student version). The unpaired two-tailed Student’s t-test was used for single-variable comparisons. Two-way repeated ANOVA followed by Sidak’s multiple comparisons were used for repeated measurements in group comparisons. Two-way non-repeated ANOVA followed by Sidak’s multiple comparisons was used for group comparisons. Error bars in graphs were represented as mean ± s.e.m. p < 0.05 was considered significant. p_∗_ < 0.05, p_∗∗_ < 0.01, p_∗∗∗_ < 0.005, p_∗∗∗∗_ < 0.0001. The sample sizes were chosen based on previously published work. All tests met assumptions for normal distribution, with similar variance between groups that were statistically compared. N values represent the final number of animals used in experiments following genotype verification and post-hoc validation of injection sites/cannula implantations.

## Acknowledgements

The authors thank Optogenetics and Viral Vectors Core supported by NIH IDDRC grant 1 U54 HD083092 and Gene Vector Core at Baylor College of Medicine for providing viral preparations. This study was supported by the NIH R21MH133228 and R01MH139750 (Y.X.). Y.X. is the awardee of the the Brain & Behavior Research Foundation awarded the first Young Investigator Grant.This study was also supported by the NIH R01DK135212, DK131446, DK136284, DK120858, DK109934, and DOD HT94252310156 (Q.T.), and NIH R01MH131570 and MH120136 (F.DM.). Q.T. is the holder of the Hans J. Muller-Eberhard, M.D., Ph.D. and Irma Gigli, M.D. Distinguished Chair in Immunology and Cullen Chair in Molecular Medicine at McGovern Medical School. Y.C. is the awardee of the Russell and Diana Hawkins Family Foundation Discovery Fellowship.

