## supplemental figures for "A Hypothalamic Inhibitory Circuit Encoding the Scalability of Stress Responses"

### Supplementary Fig. 1

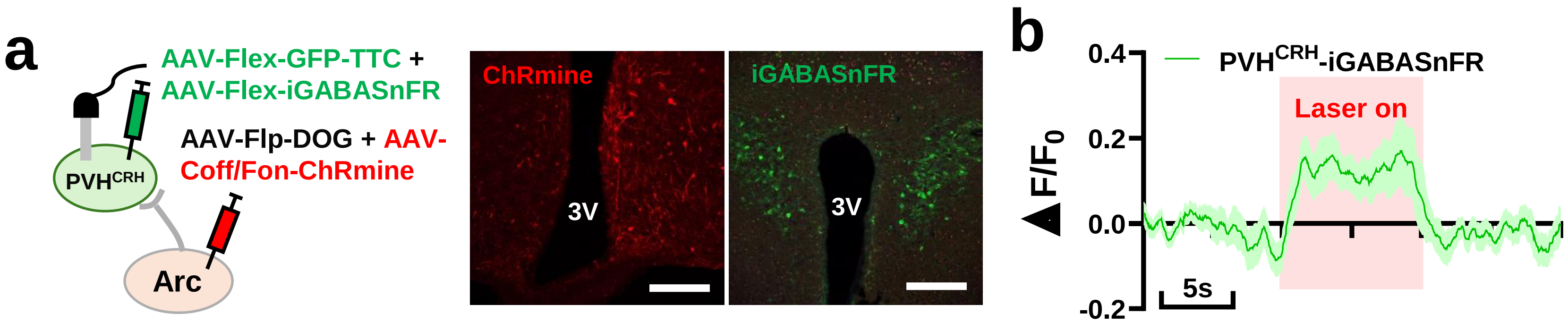


**Supplementary Fig. 1: GABA release onto PVH$^{CRH}$ neurons in response to activation of Arc$^{GABA}$ neurons. (a)** Surgical strategy and representative immunofluorescent staining images show expression pattern of iGABASnFR in PVH$^{CRH}$ neurons and ChRmine in PVH$^{CRH}$-projecting Arc neurons. (b) GABA release onto PVH$^{CRH}$ neurons show a time-locked increase pattern in response to photostimulation of PVH$^{CRH}$ -projecting Arc neurons (n = 6 mice). 3V: the third ventricle. Scar bars: 50µm.

### Supplementary Fig. 2

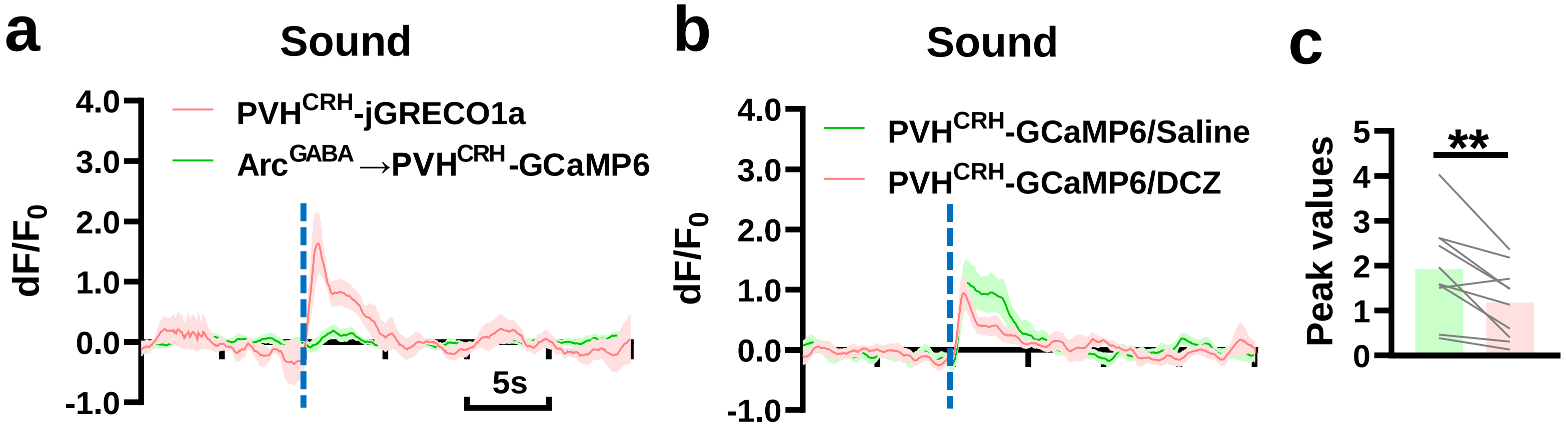


**Supplementary Fig. 2: Activity dynamics of PVH$^{CRH}$ neurons and PVH$^{CRH}$-projecting Arc$^{GABA}$ neurons during stress response induced by loud sound. (a)** In response to sound stress, PVH$^{CRH}$-projecting Arc$^{GABA}$ neurons do not show activity changes while PVH$^{CRH}$ neurons show a time-locked activity increase. **(b & c)** Chemogenetic silencing of PVH$^{CRH}$-projecting Arc$^{GABA}$ neurons resulted in a blunted activity increase of PVH$^{CRH}$ neuron activity in response to sound stressor. ** $P < 0.01$, paired *t*-test.

### Supplementary Fig. 3

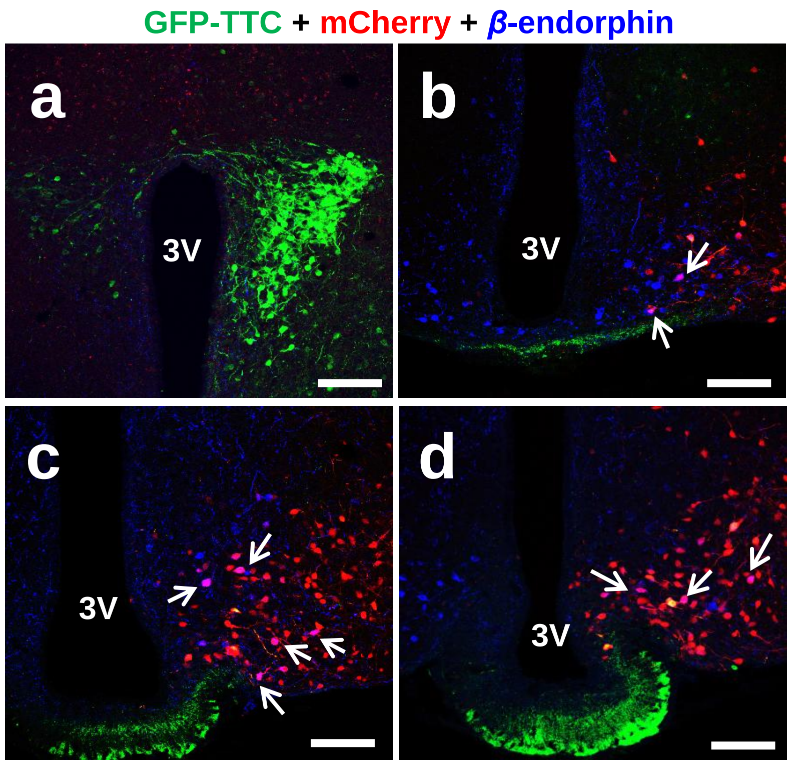


**Supplementary Fig. 3: A small portion of PVH$^{CRH}$-projecting Arc$^{GABA}$ neurons are *β*-endorphin positive.** Representative immunofluorescent staining images show that a small subset of *mCherry*-expressing Arc neurons (**b-d**, arrows) retrogradely labelled from PVH$^{CRH}$ neurons (**a**, green) co-express *β*-endorphin (**b-d**, blue), a primary marker for POMC neurons. 3V: the third ventricle. Scar bars: 50μm.

### Supplementary Fig. 4

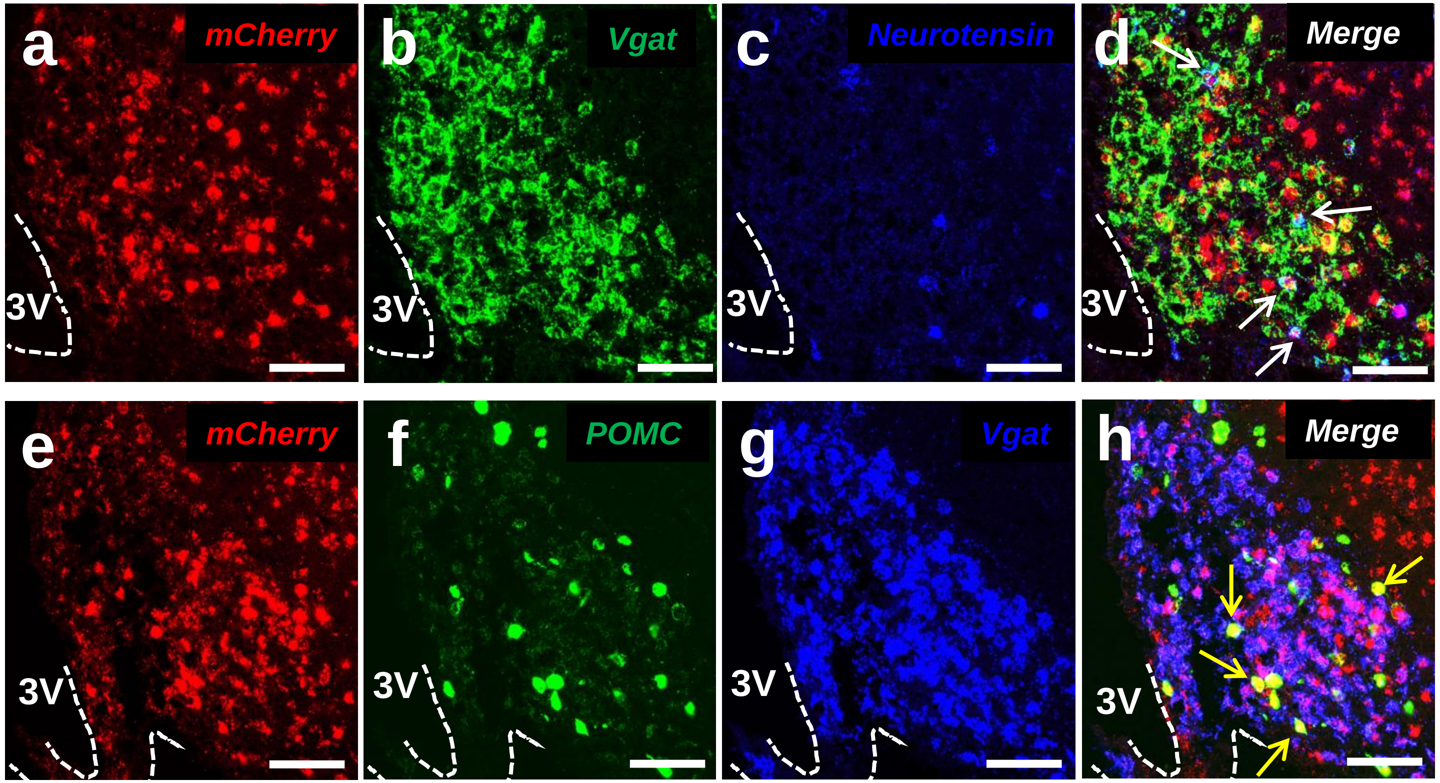


**Supplementary Fig. 4: A small portion of PVH$^{CRH}$-projecting Arc$^{GABA}$ neurons co-express *neurotensin* and *POMC*.** Representative RNAScope ISH images show that a small subset of *mCherry*-expressing Arc neurons (**d** and **h**, arrows) retrogradely labelled from PVH$^{CRH}$ neurons co-express *neurotensin* (**c**, blue) or *POMC* (**f**, green) . 3V: the third ventricle. Scar bars: 20µm.
